## Supplementary Figures for "Theoretical Morphology of Avian Wing Planform Reveals Variable Optimisation to Flight Style"

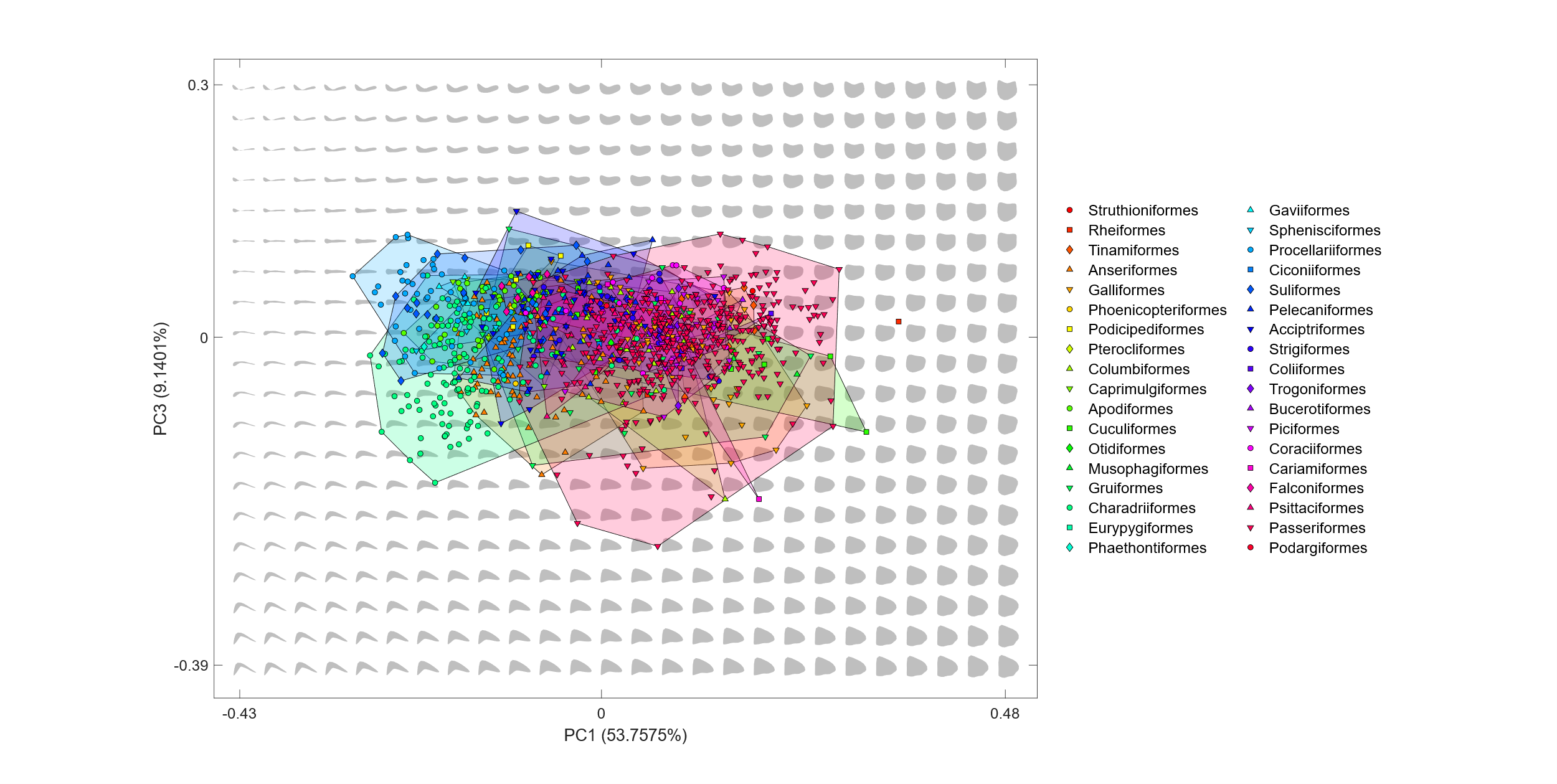

**Figure S1.** Theoretical morphospace of PC1 and PC3 with taxa separated by order. Coloured shapes represent positioning of empirical wing shapes separated at order level.

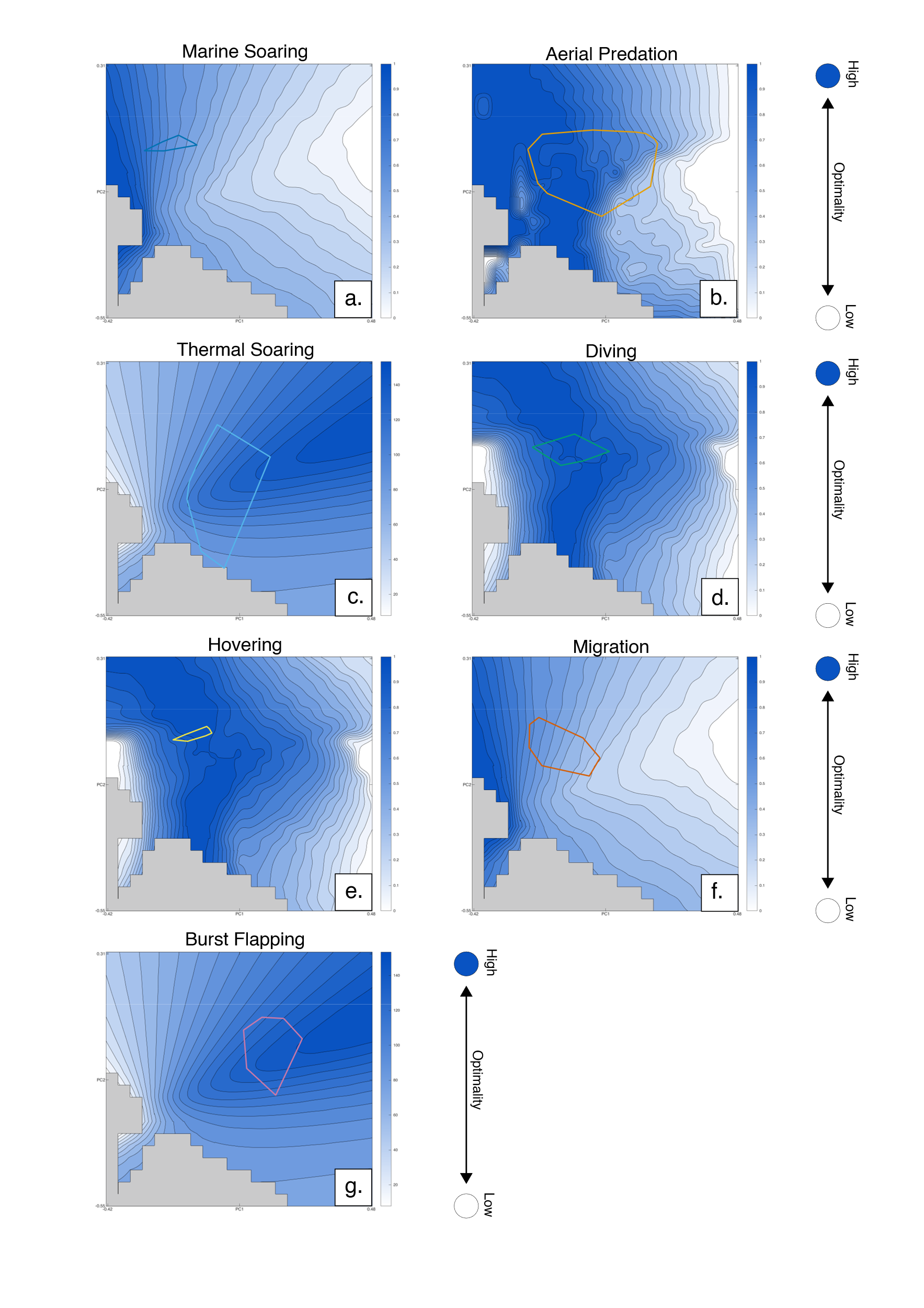

**Figure S2.** Convex hulls for specific flight styles overlaid onto the relevant performance surface for the given flight style. (**A**) marine soaring; (**B**) aerial predation; (**C**) thermal soaring; (**D**) diving; (**E**) hovering; (**F**) long-distance migration; (**G**) burst flapping.
